# The Evolution of Mass Cell Suicide in Bacterial Warfare

**DOI:** 10.1101/2020.02.25.959577

**Authors:** Elisa T. Granato, Kevin R. Foster

## Abstract

Behaviours that reliably cause the death of an actor are typically strongly disfavoured by natural selection, and yet many bacteria undergo cell lysis to release anti-competitor toxins [1–4]. This behaviour is most easily explained if only a few cells die to release toxins and help their clonemates, but the number of cells that actually lyse during bacterial warfare is unknown. The challenge is that one cannot distinguish cells that have undergone programmed suicide from those that were simply killed by a competitor’s toxin. We developed a two-colour fluorescence reporter assay in *Escherichia coli* to overcome this problem. Surprisingly, this revealed conditions where nearly all cells undergo programmed lysis. Adding a DNA-damaging toxin (DNase colicin) to a focal strain causes it to engage in mass cell suicide where around 85% of cells lyse to release their own toxin. Time-lapse 3D confocal microscopy revealed that self-lysis occurs at even higher frequencies (~94%) at the interface between competing colonies. We sought to understand how such high levels of cell suicide could be favoured by natural selection. Exposing *E. coli* that do not perform lysis to the DNase colicin revealed that mass lysis only occurs when cells are going to die anyway from toxin exposure. From an evolutionary perspective, this renders the behaviour cost-free as these cells have zero reproductive potential. This explains how mass cell suicide can evolve, as any small benefit to surviving clonemates can lead to the strategy being favoured by natural selection. Our findings have strong parallels to the suicidal attacks of social insects [5–8], which are also performed by individuals with low reproductive potential, suggesting convergent evolution in these very different organisms.

**HIGHLIGHTS:**

- A novel assay can detect *Escherichia coli* undergoing cell suicide to release toxins
- We quantified the frequency of suicidal self-lysis during competitions
- Under some conditions, nearly all cells will self-lyse to release toxins
- Self-lysis makes evolutionary sense as cells will die anyway from competitors’ toxins

## RESULTS AND DISCUSSION

*E. coli* and a number of other bacterial species release large protein toxins, often known as bacteriocins, from their cells via cell lysis [1–4]. To study the extent of cell suicide during bacterial competition with these toxins, we focussed on the well-studied group A colicins in *E. coli*. These are expressed from small, medium-copy plasmids and function to kill closely related strains and species [1]. Producing cells permeabilize their own membrane with a dedicated lysis protein to release toxins into the environment, killing themselves in the process [1–3]. When a colicin producing strain is growing alone, the colicin operon is typically only expressed in a small fraction of the population [9–15]. Expression can be upregulated by DNA damage as the colicin operon is regulated by the SOS response pathway [14,16,17], which is often done artificially via the addition of DNA-damaging agents such as mitomycin C [12,14,16,18,19]. However, many natural colicins also damage DNA, and these too have been shown to upregulate colicin production in targeted cells [9,20], an example of competition sensing [21] (Figure 1). This raises the possibility that, in competition between *E. coli* strains using DNA-damaging colicins, there may be high levels of suicidal colicin release in the population. However, to assess this, one needs to be able to distinguish between cells that are undergoing cell suicide in response to sensing a competitor’s DNA-damaging toxin, and those that were killed by the toxin itself. Moreover, we reasoned that high frequencies of a lethal behaviour would be both surprising and interesting from an evolutionary standpoint, and one that has few known precedents in the natural world. We therefore sought to develop a method that would allow us to make the critical distinction between a cell killing itself via an evolved behaviour, and one simply dying due to exposure to a DNA-damaging agent.

**Figure 1.**
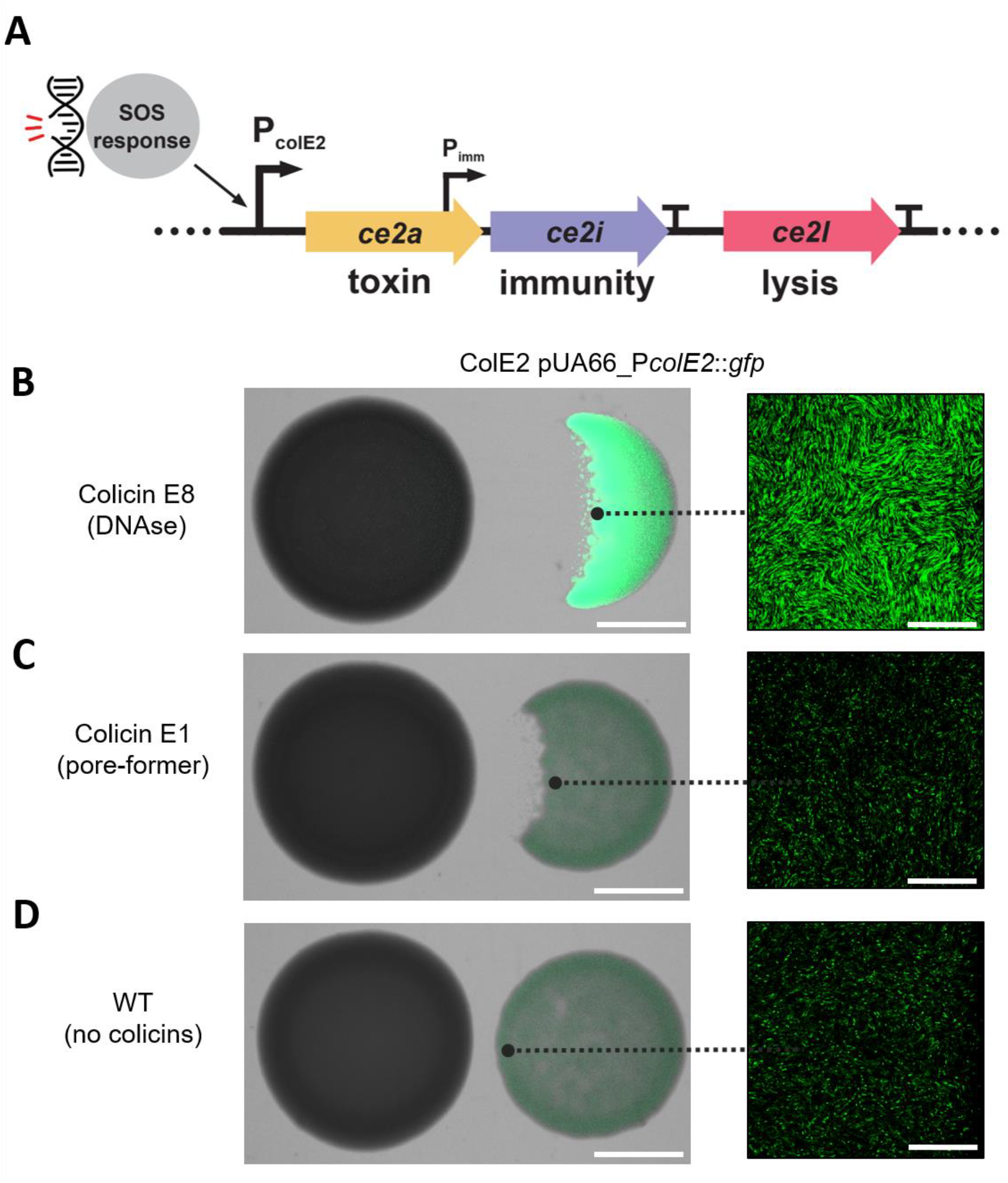
*E. coli* activates the colicin operon in response to a competitor causing DNA damage. **(A)** Overview of the colicin operon as encoded on plasmid pColE2-P9 in *E. coli*. In response to DNA damage, the P_colE2_ promoter is activated and the genes encoding colicin E2 (*ce2a*) and its cognate imunity protein (*ce2i*) are expressed. To ensure sufficient levels of immunity protein, *ce2i* is additionally transcribed from a second, constitutively active promoter (P_imm_) located within the *ce2a* gene. A transcriptional terminator (T) after the immunity gene *ce2i* ensures that the downstream gene encoding the lysis protein (*ce2l*) is only expressed at very high levels of P_colE2_ activation. Adapted from [1]. **(B)** The response of the *E. coli* colicin E2 promoter to a foreign DNase colicin (colicin E8). *E. coli* BZB1011 colonies producing colicin E2 were grown next to competitor colonies overnight and then imaged by stereomicroscopy (left) and confocal microscopy (right). In the colicin E2 producer, the colicin E2 promoter also drives the expression of GFP on a reporter plasmid (pUA66-P*colE2*::*gfp*). **(C)** Absence of response of the *E. coli* colicin E2 promoter to the pore-forming colicin E1 made by *E. coli* BZB1011 carrying the colicin E1 plasmid. **(D)** Wildtype control where the lefthand strain (BZB1011) lacks any colicin plasmid and so does not produce colicins. Scale bars: 2 mm (left), 50 µm (right).

### Self-lysis frequency is modulated by competitor toxin concentrations

We followed the production of the colicin E2 toxin and its cognate immunity protein (Figure 1A) using a reporter plasmid that expresses green fluorescent protein (*gfp*) from the native colicin E2 promoter [9] (pUA66-PcolE2::*gfp*, Figure 1B-D). This construct, like others previously studied [10,12,22,23], allows one to identify cells that may be on their way to cell suicide via lysis [1–3]. However, it is not sufficient to follow the full behaviour, as the self-lysis process is dependent on the expression of the lysis gene in the colicin operon [18,24], which is subject to several additional layers of regulation at both the transcriptional and post-transcriptional level [17,25–28]. Moreover, with this reporter alone, it is impossible to tell if a cell that died was killed by external toxins while building up colicins or if it had undergone self-lysis. We therefore sought to identify ways to distinguish the self-lysis phenotype from cell death and discovered that one of the standard DNA dyes used in microbiology – propidium iodide – reliably identified cells that have undergone self-lysis. Propidium iodide (hereafter PI) is typically used as a ‘death’ stain, because it will only enter cells with a compromised membrane and bind their DNA, generating a fluorescent signal [29]. We found that self-lysis leads to PI reliably entering cells to give the fluorescent signal but, critically, cell death caused by the DNase colicin toxin of another *E. coli* strain does not. When combined with the fluorescent signal from the GFP reporter plasmid [9], this results in a characteristic two-colour fluorescent pattern when cells are undergoing self-lysis, but not when they are killed by foreign DNase toxins (Figures 2A and 2B, Suppl. Figure S1, Suppl. Movies S1 and S2).

**Figure 2.**
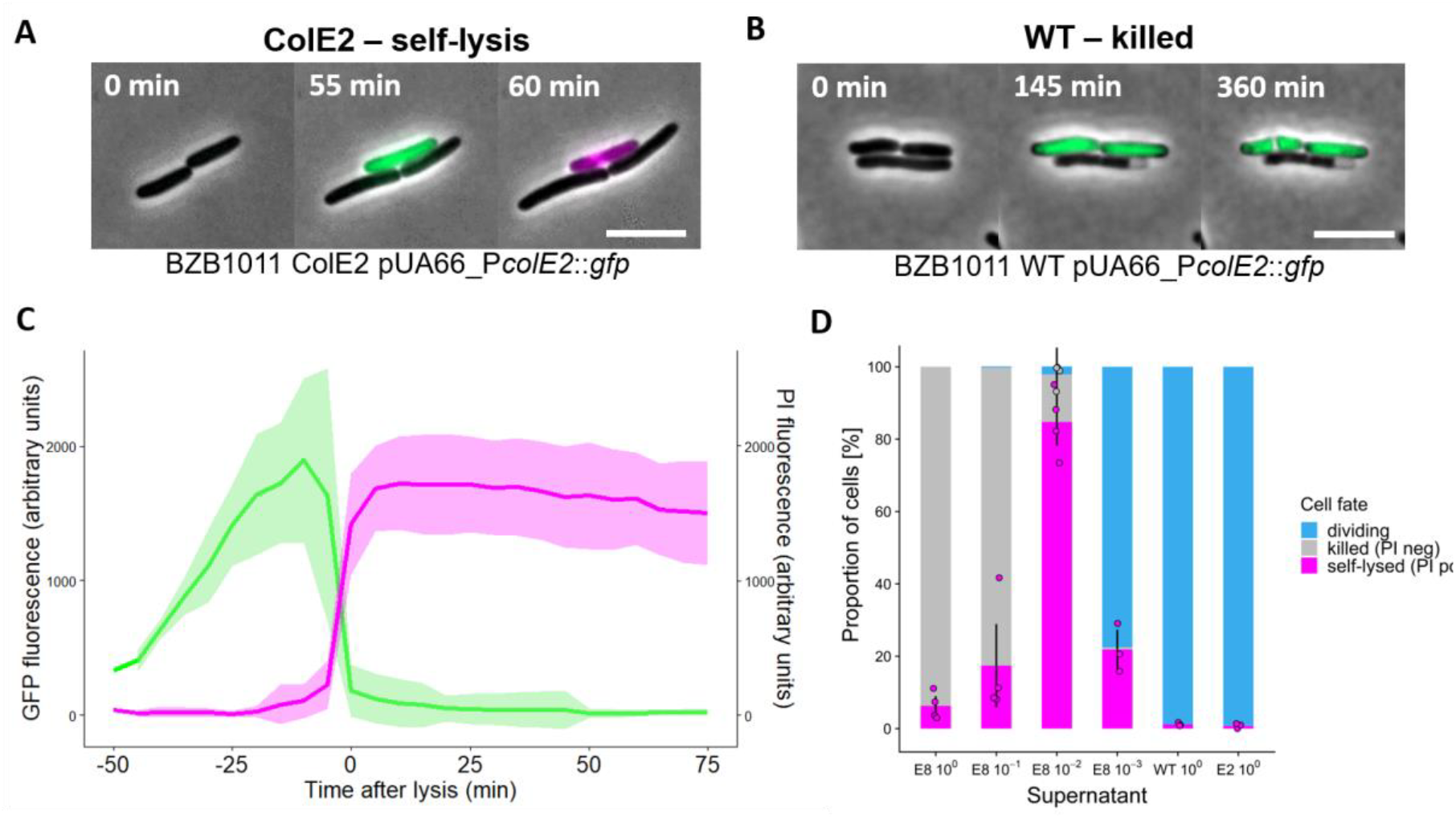
Self-lysis frequency in E. coli is modulated by competitor toxin concentrations. (**A)** Representative time-lapse micrographs of ColE2 pUA66-P*colE2*::*gfp* cells undergoing self-lysis. Phase-contrast channel, GFP channel and propidium iodide (PI) channel are overlaid in all images. GFP signal indicates colicin promoter activation. PI signal indicates membrane permeabilization, i.e. self-lysis. Scale bar, 5 µm. See Movie S1. (**B)** Representative time-lapse micrographs of wildtype cells (WT pUA66-P*colE2*::*gfp*) being killed by a foreign DNase colicin. Phase-contrast channel, GFP channel and propidium iodide (PI) channel are overlaid in all images. The absence of PI signal in a non-dividing, dead cell is indicative of an intact membrane and killing by the action of the foreign DNase colicin. Scale bar, 5 µm. See Movie S2. (**C)** Fluorescence signals in cells undergoing self-lysis in response to colicin E8. ColE2 pUA66-P*colE2*::*gfp* cells were exposed to a 1% dilution of supernatant of a colicin E8-producing strain and imaged for up to six hours. Individual cell fluorescence tracks are shown for the GFP channel (green) and PI channel (magenta). Thick lines and shaded areas indicate the mean and standard deviation across n = 20 lysed cells in the same field of view. See Movie S3. Figure S1 and Movie S4 show a negative control where colicin E8 is added to wildtype cells that lack the colicin plasmid. (**D)** Cell fate frequencies in populations of *E. coli* exposed to colicin E8. ColE2 pUA66-P*colE2*::*gfp* cells were exposed to different dilutions of supernatant of a colicin E8-producing strain, a non-producing wild-type, or their own sterile supernatant. Cells were imaged for up to six hours, and the fate of n = 7985 cells across all treatments was categorized as either dividing, killed (non-dividing and PI-negative) or self-lysed (non-diving and PI-positive). A Kruskal-Wallis test yielded a statistically significant relationship between supernatant concentrations and self-lysis frequencies (Chi square = 18.285, df = 5, p = .0003).

Using this assay, we set out to characterize the response of a focal colicinogenic strain to a competitor producing a foreign DNA-damaging colicin (E8), which is expected to elicit SOS response, colicin production and subsequent self-lysis. We began by exposing the colicin E2-producing strain (ColE2) carrying the reporter plasmid (pUA66-PcolE2::*gfp*) to sterile supernatants of a colicin E8-producing competitor on nutrient agar containing PI. Using time lapse fluorescence microscopy and cell tracking, we measured GFP and PI fluorescence levels in thousands (n = 7985) of individual cells exposed to colicin E8 at a range of concentrations (Figures 2C and 2D, Suppl. Movie S3). In cells undergoing self-lysis, the GFP signal steadily increased for approximately 50 minutes leading up to the lysis event, representative of the continuous activation of the colicin promoter in response to the imposed DNA damage, and thus accumulation of colicins in the cytoplasm (Figure 2C). Self-lysis events were then characterized by the simultaneous influx of PI binding to DNA (increased red fluorescence signal) and efflux of GFP out of the cell (decreasing GFP signal) due to leakage of the cytoplasm into the environment (Figure 2C).

We combined these fluorescence measurements with cell viability observations to divide individual cell’s responses to colicin E8 into three categories (Figure 2D): non-dividing and PI-negative (killed cell) or PI-positive (self-lysed cell), as well as dividing and PI-negative (live cell). At the highest concentration of toxic supernatant, most cells were immediately killed upon exposure to colicin E8, and little self-lysis occurred (6.3%). In treatments with successively lower concentrations of colicin E8 however, the proportion of self-lysed cells increased dramatically as cells had more time to respond to the stress, reaching a peak at 1% supernatant concentration of 84.7% self-lysed cells on average, with a maximum value of up to 95% in one experiment (Figure 2D). At lower concentrations of toxin-containing supernatant, the self-lysis response became less frequent and more cells started to divide (21.9% self-lysis), which is expected as the amount of DNA damage experienced by the cells should be much lower. Taken together, these results show that colicinogenic cells respond to DNA-damaging competitor colicins with self-lysis in a concentration-dependent manner. Moreover, we find conditions where cells will undergo mass self-lysis with the great majority of cells dying to relase toxins.

### Mass cell suicide is also seen in competitions between *E. coli* strains

Our first experiments show that adding supernatant containing a DNase colicin results in a dose-dependent self-lysis response, with conditions where nearly all cells lyse. However, in nature, bacteria are likely to be exposed to toxin concentrations that change over both time and space as they compete with adjacent populations of cells [30,31]. To capture these effects, we followed the lysis response in a monolayer colony of our focal strain (ColE2 pUA66-PcolE2::*gfp*) with a colicin E8-producing competitor growing next to it (Suppl. Movie S5). In addition to allowing for toxin concentrations to change in time and space, this setup also allows the two strains to react to each other. In particular, both strains use DNase colicins that will activate the SOS response and trigger colicin production in their, an example of competition sensing [9,21,32]. We seeded the colicin E2-producing strain carrying the reporter plasmid (ColE2 pUA66-PcolE2::*gfp*) or a non-producing wildtype strain (WT pUA66-PcolE2::*gfp*) in thin monolayer colonies on nutrient agar, with their colicin E8-producing competitor growing next to them. Using time lapse fluorescence microscopy at the interface between the two colonies, we can then track thousands (n = 3420) of individual cells as they react to the incoming flow of colicin E8 toxins (Suppl. Movie S5), and measure self-lysis frequencies as a function of their distance from the interface (Figure 3). As expected, in the wildtype control we observed that very few cells exhibited a PI fluorescence signal, consistent with PI being a good indicator of self-lysing cells. For the colicin E2-producer, we found that self-lysis frequencies within six hours of observation were on average 75.5% at the very colony edge, and 97.3% in cells positioned slightly further (ca. 0.5 mm) away from the edge. For cells positioned even further away from the edge (> 1.2 mm), self-lysis frequencies rapidly decreased to 0.8% (Figure 3B).

**Figure 3.**
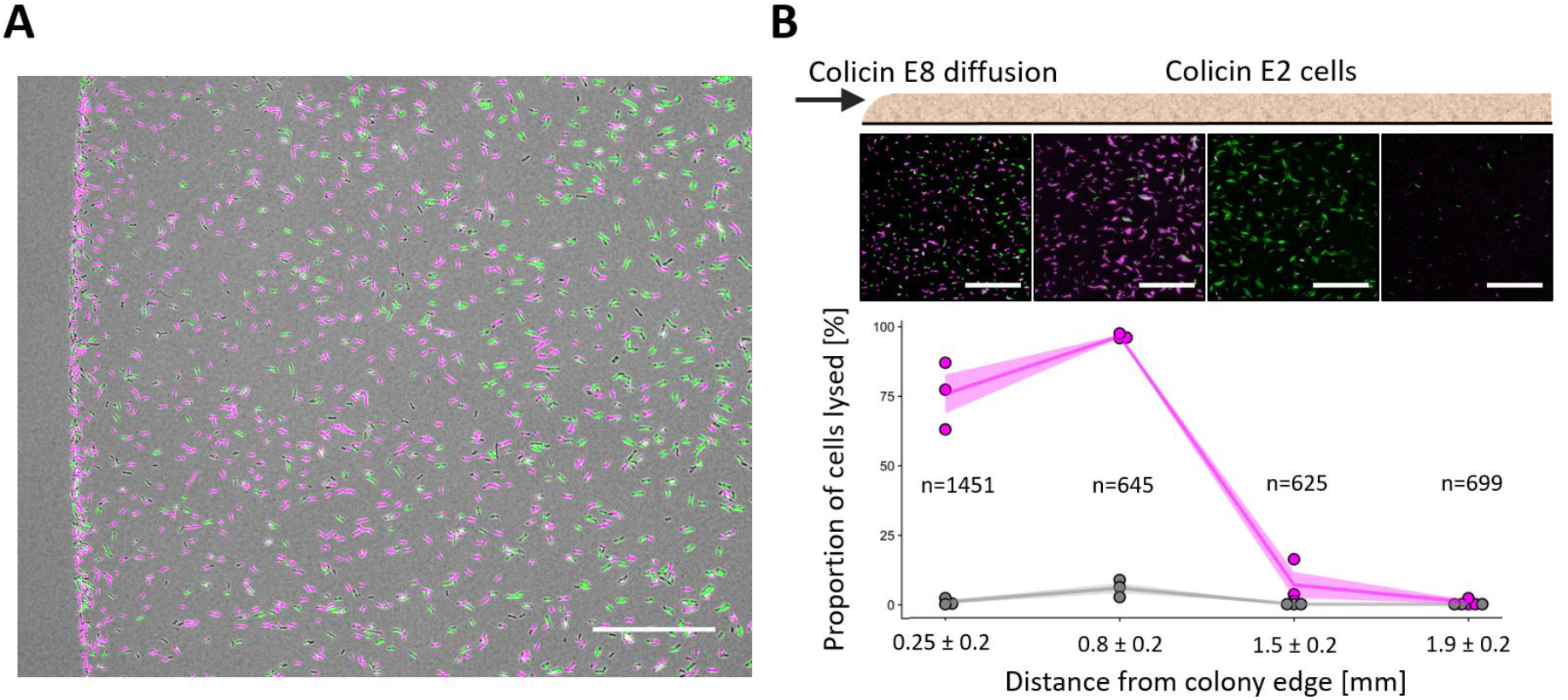
Cells facing a competitor exhibit local mass self-lysis. Self-lysis quantification in thin colonies exposed to colicin E8 produced by a nearby colony. Cells of a focal strain either capable of producing colicin E2 and self-lysing (ColE2 pUA66-P*colE2*::*gfp*) or a wildtype non-producer incapable of self-lysis (WT pUA66-P*colE2*::*gfp*) were seeded in a monolayer colony on nutrient medium supplement with propidium iodide (PI, 1 μg/mL) next to a strain producing colicin E8. Cells of the focal strain were imaged using time-lapse fluorescence microscopy at four locations in the colony situated at different distances from the colony edge facing the competitor. **(A)** Representative snapshot of ColE2 pUA66-P*colE2*::*gfp* colony edge after two hours of exposure to a ColE8 competitor colony (not in view), showing self-lysed cells in magenta (PI signal) and colicin promoter activity in green (GFP signal). Scale bar, 100 μm. **(B)** Individual cells of strain ColE2 pUA66-P*colE2*::*gfp* (magenta) or WT pUA66-P*colE2*::*gfp* (grey) were tracked over six hours (n = total numbers of cells across both strains successfully tracked over ≥ four hours), and self-lysis frequencies were quantified by measuring PI-specific fluorescence in individual cells and determining the frequency of PI-positive tracks relative to the total number of cells tracked in that position. Lines and shaded areas indicate the mean and SEM across three biological replicates. Images above each distance point show representative fluorescence images of one replicate colony acquired at the respective position after four hours of observation. Scale bar, 100 μm. The image shown in (A) covers the same area as the leftmost of the fluorescence images shown in (B). We detected a statistically significant difference between the two strains in their frequency of PI-specific fluorescence at distances 0.25 and 0.8 mm (linear model: F(1,4) = 114.1 for distance = 0.25, 2368 for distance = 0.8; p < .001). See Movie S5.

This strong spatial relationship is likely explained by the competitor toxin diffusing towards and through the colony, leading to different micro-environments for cells in different locations. Consistent with this, at the very colony edge, the cells that did not lyse failed to divide (Suppl. Movie S5), indicating that these cells experienced lethal stress levels that killed them so quickly that it prevented the self-lysis response. However, cells further into the colony were able to achieve mass self-lysis, presumably because the toxin built up more slowly at their more distant position, preventing immediate death and allowing for widespread self-lysis. As expected from this interpretation, we see more and more cell division at locations further into the colony, as cells get more time to grow and divide before being exposed to the toxin. At this point, stress levels in a large fraction of the population failed to reach the critical threshold required for induction of the lysis gene, resulting in a strong decline in self-lysis frequency (Figure 3B). These results demonstrate that when colicinogenic colonies are exposed to a competitor secreting a DNase toxin, self-lysis frequencies strongly depend on the distance from the competitor, and peak at intermediate distances, where cells are not immediately killed but still experience high enough stress levels to trigger self-lysis. They also suggest that the extremely high self-lysis frequencies observed in response to supernatants (Figure 2D) represent realistic behavioural patterns that can be recapitulated in spatially structured colonies (Figure 3B).

Bacteria often grow in dense biofilms or host-associated communities, where cell-cell interactions between clonemates - in addition to interactions between genotypes - can play a key role in their behaviours [30,33–35]. In the case of colicins, the production of some DNase colicins - including colicin E2 - can be induced by the presence of the same colicins released by clonemates, a phenomenon termed *autoinduction* [9,15,36]. We therefore sought to follow the lysis response in bacterial colonies at high cell density, where autoinduction is expected to be maximal. To image single cell behaviour at the interface between two dense bacterial communities, we used time lapse 3D confocal microscopy, which allows for single cell resolution of fluorescence signals over space and time. We followed a dense multilayer colony of colicin E2-producing cells constitutively producing GFP (ColE2 *gfp*), with a colicin E8-producing strain growing next to it (Suppl. Movie S6). This allowed us to measure self-lysis frequencies as percent of total biomass over both time and as a function of their distance from the colony edge facing the competitor (Figure 4). We found that within eight hours of observation 60.1%-75.8% of the total biomass self-lysed at the very edge of the colony (< 0.1 mm), and maximum self-lysis levels of 93.5%-96.4% was reached in locations further away from the edge (0.2 mm – 0.4 mm). For cells positioned even further away from the edge (> 0.5 mm), self-lysis frequencies decreased again to 55.8%-77.0% (Figure 4B). This spatial dependence of self-lysis frequencies mirrors our observations in thin monolayer colonies (Figure 3B), and is likely caused by some cells at the very edge dying before self-lysing, while cells further into the colony exhibit near-complete self-lysis. These results demonstrate that mass self-lysis also occurs in three-dimensional, high-density bacterial colonies in response to a competitor.

**Figure 4.**
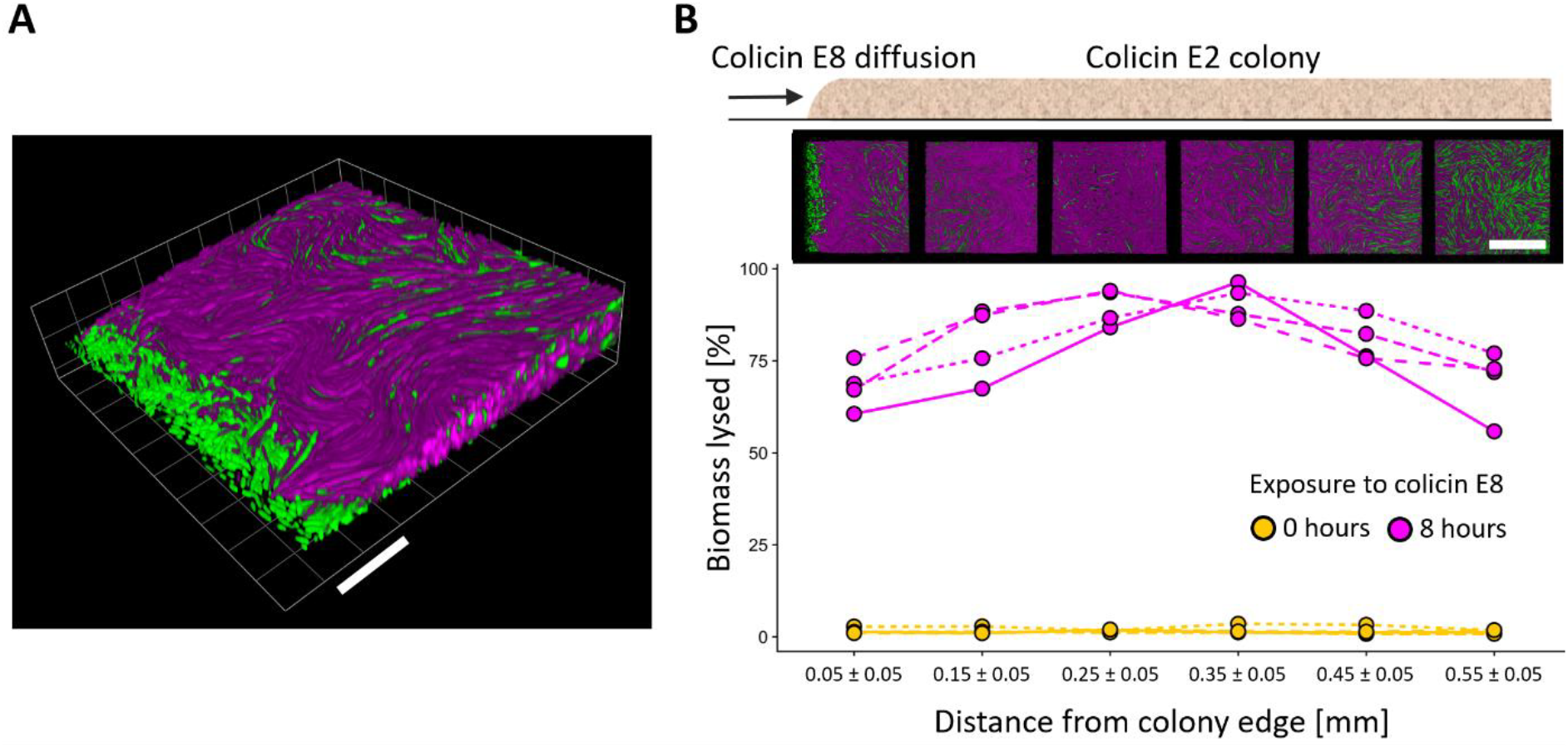
Dense colonies facing a competitor exhibit local mass self-lysis. Self-lysis quantification in three-dimensional colonies exposed to colicin E8 produced by a nearby colony. Cells of a focal strain producing GFP constitutively and capable of producing colicin E2 and self-lysing (ColE2 *gfp*) were grown in a three-dimensional colony on nutrient medium supplement with propidium iodide (PI, 1μg/mL) next to a strain producing colicin E8. The focal colony was imaged for eight hours using time-lapse 3D confocal microscopy at six locations situated at different distances from the colony edge facing the competitor. **(A)** 3D rendering of confocal image of a colony edge after eight hours of exposure to colicin E8. Scale bar, 20 μm. **(B)** Self-lysis at each location was quantified by determining the volume of biomass exhibiting PI-specific fluorescence (indicating self-lysis) relative to the total biomass after zero or eight hours of exposure to colicin E8. Total biomass was calculated by determining the volume of biomass exhibiting GFP-specific fluorescence (indicating either live cells or dead cells that did not self-lyse) plus biomass of those exhibiting PI-specific fluorescence. Line-types indicate four independent biological replicates. We detected a statistically significant increase in PI-specific fluorescence over the observation period at all locations (linear model: *F*(1,6) > 212.2; p < .001). Images above each distance point show 3D renderings of confocal images of the same replicate colony acquired at the respective position after eight hours of observation, viewed from above. Scale bar 50 μm. See Movie S6. See Figure S2 for the same experiment using a wildtype, non-producing control as the focal strain.

### Mass cell suicide is associated with the loss of reproductive potential

Our discovery of massive cell suicide in bacteria, to the extent that locally more than 95% of cells will lyse themselves, is surprising given the typically strong natural selection against such individually-costly behaviours. One important component to the explanation is that clonemates must benefit from the behaviour and maintain the trait in the population (an example of kin selection) [37–39]. Consistent with this, we see in colony experiments that only the front line of cells engage in mass cell suicide with the potential of clonemates behind to benefit from their behaviour (Figures 3 and 4). However, even with this explanation, it remains surprising that so many cells engage in the behaviour, which removes nearly all cells from the edge of the colony and greatly slows expansion into new territory. To understand the conditions of self-lysis better, therefore, we asked what would have happened to these cells had they not been able to self-lyse. We repeated the supernatant exposure experiment (Figure 2D) with a control strain that is genetically identical to the self-lysing strain except that it lacks the colicin plasmid, and thus the ability to produce colicins and self-lyse (WT pUA66-PcolE2::*gfp*). This strain still carries the reporter plasmid (Figure 2B), which indicates whether the colicin promoter would be active if the cells were able to make colicins, and it can be targeted and killed by colicin E8. We exposed this strain to supernatants containing colicin E8 at a range of concentrations and quantified the fate of the exposed cells (Figure 5A). Importantly, under conditions where we observed the highest levels of self-lysis in the colicinogenic strain (Figure 2D, third bar), we observed that the great majority of wildtype control cells were dead (Figure 5A, third bar). This indicates that the level of DNA damage these cells experienced when exposed to the DNase colicin E8 was too much to be repaired, resulting in their death. Therefore, the colicinogenic cells that self-lysed *en masse* under the same conditions (Figure 2D) most likely experienced a lethal level of stress imposed by the competitor toxin. The fitness costs of the response are thus mitigated by lysing cells having low or no future reproductive potential.

**Figure 5.**
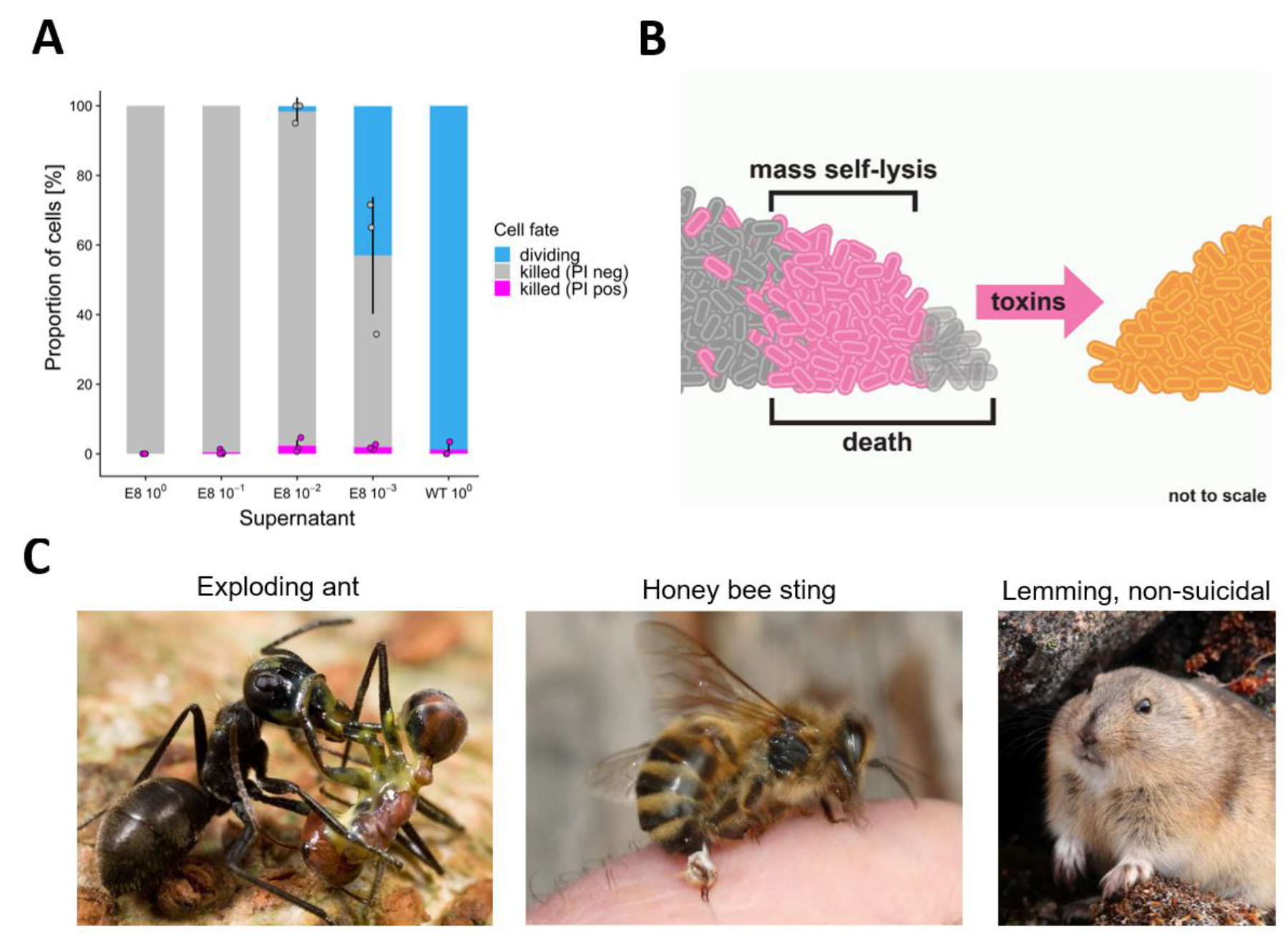
Suicidal behaviour is associated with low reproductive potential. **(A)** Cell fate frequencies in response to colicin E8 in *E. coli* cells unable to self-lyse. WT pUA66-P*colE2*::*gfp* cells were exposed to different dilutions of supernatant of a colicin E8-producing strain or a non-producing wildtype. Cells were imaged for up to six hours, and the fate of n = 3666 cells across all treatments was categorized as either dividing, dead and PI-negative, or dead and PI-positive. A Kruskal-Wallis test did not yield a statistically significant relationship between supernatant concentrations and frequency of cells positive for PI-specific fluorescence (Chi square = 7.8486, df = 4, p = .09). **(B)** Illustration of the mass cell suicide phenotype. In the region where the toxins of the competing strain (orange) reaches lethal levels, large number of cells of the focal strain lyse (pink) and release colicins en masse (pink arrow). Cell suicide does not occur at the very edge of the colony on the left (transparent grey cells), as these cells die immediately from the competitors toxins. **(C)** Two examples of suicidal behaviors in social insects, and a popular misconception. Left: A minor *Colobopsis cylindricus* (“exploding ant”) worker has ruptured her body to release a sticky yellow substance, killing both herself and her opponent, the larger worker of another ant species (*Camponotus sp.*) [6,55]. Middle: Honey bee (*Apis mellifera*) workers sting in defense of their colony, which often results in the worker’s death as the sting gets pulled out of her body [7]. Right: A long-standing myth incorrectly holds that lemmings commit mass suicide. The example is illustrative because - unlike in the social insects and bacteria where low reproductive potential and benefits to kin can explain suicidal behaviours - there is no evolutionary rationale for such behaviour in lemmings. Image sources: exploding ant (Mark W. Moffett, used with permission); honeybee (credit: Waugsberg/Wikimedia Commons (CC BY-SA 3.0)); lemming (credit: Argus fin/Wikimedia Commons (public domain)).

## CONCLUSIONS

We have identified competition scenarios where bacteria will engage in mass cell suicide, with the proportion of cells engaging in the behaviour reaching almost 100% in some areas (Figure 5B). It is important to acknowledge that not all bacteria display this behaviour: the suicidal release of toxins is known from several bacterial species [1,4] but it is not the norm across species. Moreover, in the case of *E. coli*, we only expect these levels of cell lysis to occur when a focal strain is targeted by DNA damaging toxins that allow it to detect the incoming attack and respond with its own colicin. Some colicins have other mechanisms of action, including pore-forming colicins that are ‘silent’ toxins that can kill cells without activating the SOS response or toxin release in a competitor [9,32] (Figure 1C). Nevertheless, the high frequency of cell suicide we see when DNase producing strains meet is striking and represents one of the most extreme social phenotypes documented to date in bacteria or elsewhere. A twist in the case of colicin production is that the genes for the behaviour are carried on plasmids that, under some conditions, can be subject to horizontal transfer between strains via conjugation [40]. This has the potential to lead to additional complexities where natural selection on the plasmids differs from that of their hosts. However, colicin plasmids cannot always conjugate (as in our experiments) and suicidal cell behaviours are encoded in the chromosome in species like *Pseudomonas aeruginosa* [4]. The potential for conjugation, therefore, is not a requirement for the evolution of toxin release via cell lysis.

The evolution of self-lysis is also known from the phenomenon of abortive phage infection in bacteria [41–43]. Here, infected cells have evolved to lyse rather than pass on the infection to clonemates, and the great majority of cells will undergo lysis if enough phage are added to a culture [44]. However, in terms of function, much more similar are examples from the social insects. Massive suicidal attacks are known from several insect species (Figure 5C) where they are performed by the old workers that have the lowest reproductive and helping potential in the colony [5–8]. We see a clear parallel in our experiments where the highest levels of cell suicide occur under conditions where the cells are going to die anyway from exposure to the toxin. These cells then are no longer able to divide and, as in the social insect examples, this loss of reproductive potential is associated with the expression of the attack behaviour. This makes the fitness costs associated with this extreme behaviour close to zero and, so long as there are some clonemates in the vicinity that can benefit, the behaviour can be favoured by natural selection (more specifically kin selection [37,45]). Our work suggests a strong evolutionary convergence in the sociality of bacteria and that of the social insects. In both, examples of mass self-sacrifice are seen in defence of the colony and, in both, these behaviours can be explained by low personal fitness costs combined with benefits to kin.

## MATERIALS AND METHODS

### Bacterial growth conditions and strain construction

Unless otherwise indicated, all *E. coli* BZB1011 strains were grown overnight in 5 ml LB medium (per L: 10g Tryptone, 10g NaCl, 5g Yeast Extract) in 15 ml polypropylene tubes at 37°C with agitation (220 rpm). When necessary, the medium was supplemented with kanamycin (50 μg/mL). All experiments were carried out at 37°C. For time-lapse microscopy experiments, samples were kept at 37°C at all times using a custom-built incubation chamber. Whole-colony competitions (Figure 1B-D) were carried out on 0.8% w/v LB Agar. Supernatant exposure assays (Figures 2, 5A and S1) were carried out on 0.8% w/v LB Agarose supplemented with 1 µg/mL propidium iodide (PI; Sigma-Aldrich). Colony monolayer assays (Figure 3) and 3D colony imaging experiments (Figures 4 and S2) were carried out on 0.8% w/v LB Agar supplemented with 1 µg/mL PI. For the generation of the BZB1011 WT pUA66-P*colE2*::*gfp* strain, BZB1011 WT cells were transformed with the reporter plasmid pUA66-P*colE2*::*gfp* [9] via electroporation and selected on 50 μg/mL kanamycin at 37°C. All plasmids and strains used in this study are listed in Table S1.

### Whole-colony competitions

To monitor the response of a colicin E2 producing strain to foreign colicins with different cellular targets (Figure 1B-D), cells were grown to exponential phase, washed twice with LB medium, and resuspended and normalized in LB medium to an optical density at 600 nm (OD600) of 1.0 for the competitor (ColE8, ColE1, or WT as a negative control) and a 10^−3^ dilution of OD600 = 1.0 for the focal strain (ColE2 pUA66-P*colE2*::*gfp*). Per strain combination, three 5 μl spots of bacterial suspension were then spotted next to each other onto 0.8% w/v LB agar plates, resulting in a distance of approximately 0.5 mm between the respective spot edges. Plates were then incubated at 37°C for 12 hours. After incubation, bright field and GFPmut3-fluorescence (ex: 500 nm|em: 513 nm) images of whole colonies were acquired using a Zeiss Plan-Apochromat Z 0.5 × objective on a Zeiss AxioZoom.V16 stereomicroscope with ZEN Blue software (version 1.1.2.0). Colony edges of the same colonies were then surface-imaged using a Zeiss EC Epiplan-Neofluar 50x air objective (NA = 0.8) on a Zeiss LSM880 confocal laser scanning unit in regular confocal mode, using ZEN Black software (version 14.0.18.201). Images were analysed using *FIJI* [46], and Figure 1B-D shows representative images of three biological replicates.

### Supernatant exposure assay

To record time courses of cells reacting to sterile supernatant of a competitor, cells of the competitor strain (ColE8, ColE2, or WT as a control) were grown overnight for 16 hours, and 1 mL of bacterial suspension was centrifuged for 5 min at 17500*g to sediment cells. The resulting supernatant was sterile-filtered using a 0.2 μm syringe filter (Sartorius) and then serially diluted in sterile saline (0.8% NaCl in ddH_2_O). 3 μL of undiluted or diluted sterile supernatant were then pipetted onto small circular cut-outs (diameter 5mm, height 2mm) of 0.8% w/v LB agarose + PI and left to dry and diffuse in the agarose pad for 1 hour at room temperature. In parallel, cells of the focal strain (ColE2 pUA66-P*colE2*::*gfp*, or WT pUA66-P*colE2*::*gfp*) were grown to exponential phase (OD600 ~ 0.2), washed twice with LB medium, resuspended, and then diluted 1:50 with LB medium. 1 μl of this diluted bacterial suspension was then spotted onto the agarose pads infused with supernatant and left to dry for 10 minutes at room temperature. Pads were then placed onto a glass slide and covered with a 22×22mm n° 1.5 coverslip so that the cells were sandwiched between the agarose and the coverslip. The sides of the coverslip were sealed with *Glisseal-HV* laboratory grease (VWR) to avoid desiccation. The sample was then moved to the microscope and imaged immediately. Time-lapse fluorescence microscopy was performed using a Zeiss Axio Observer inverted microscope with a Zeiss Plan-Apochromat 63x oil immersion objective (NA = 1.4) and ZEN Blue software (version 1.1.2.0). Exposure times were 156 ms for phase contrast, 100 ms for GFPmut3 (ex: 500 nm|em: 513 nm), and 50 ms for PI (ex: 493 nm|em: 636 nm). Images were acquired every 5 minutes for 6 hours. Image analysis and cell tracking was carried out with *FIJI* [46] and *FAST* [47]. For the data presented in Figures 2D and 5A, three or four biological replicates were carried out for each supernatant concentration and strain combination, and for each replicate at least three fields of view were recorded at each time point. A total number of n = 7985 and 3266 cells were tracked for the two focal strains ColE2 pUA66-P*colE2*::*gfp* and WT pUA66-P*colE2*::*gfp*, respectively. Cells that failed to divide during the six hour observation time were counted as dead, and cells that started dividing were counted as such. Cells that exceeded a PI-specific fluorescence value over a threshold of 3x the background value during the observation period were categorized as PI-positive, cells below that value as PI-negative. The data presented in Figure 2C is a subset of the data presented in Figure 2D and shows background-corrected and GFP- and PI-specific fluorescence values for all tracked ColE2 pUA66-P*colE2*::*gfp* cells in one field of view exposed to 1% ColE8 supernatant. For visualization purposes, time-series values were synchronized with respect to time-to-lysis in each cell.

### Colony monolayer imaging

To image colony monolayers of a focal strain reacting to a nearby competitor, cells were grown to exponential phase, washed twice with LB medium, and resuspended and normalized in LB medium to an OD600 of 1.0 for the competitor (ColE8) and 0.05 for the focal strain (ColE2 pUA66-P*colE2*::*gfp*, or WT pUA66-P*colE2*::*gfp*). 5 μl of bacterial suspension per strain were then spotted next to each other onto 0.8% w/v LB agar + PI plates, resulting in a distance of approximately 0.5 mm between the spot edges. After drying for 10 minutes at room temperature, a ca. 2×2 cm section of agar around a pair of seeded spots (competitor and focal strain) was then cut out using a scalpel, and placed onto a 5 cm diameter glass bottom Petri dish with a 3-cm diameter uncoated n°1.5 glass window (MatTek Corporation), which was then inverted to allow immediate imaging of the spots through air. Time-lapse imaging of the spot edge at the interface between the competitor and the focal strain was then performed using a Zeiss Axio Observer inverted microscope with a Zeiss Plan-Apochromat 20x objective (NA = 0.8) and ZEN Blue software (version 2.6.76.00000). Exposure times were 2.72 ms for bright-field, 20 ms for GFPmut3 (ex: 500 nm|em: 513 nm), and 80 ms for PI (ex: 493 nm | em: 636 nm). Images were acquired every five minutes for six hours, and image analysis and cell tracking was carried out with *FIJI* [46] and *FAST* [47]. For the data presented in Figure 3B, three biological replicates were carried out for each strain, and for each replicate six fields of view at different distances from the colony edge were recorded at each time point. A combined total number of n = 3420 cells were successfully tracked over ≥ four hours for the two focal strains ColE2 pUA66-P*colE2*::*gfp* and WT pUA66-P*colE2*::*gfp*. Cells that exceeded a PI-specific fluorescence value over a threshold of 3x the background value during the observation period were categorized as PI-positive, cells below that value as PI-negative.

### 3D colony imaging

To image three-dimensional colonies of a focal strain reacting to a nearby competitor, cells were grown to exponential phase, washed twice with LB medium, and resuspended and normalized in LB medium to an OD600 of 1.0 for both the competitor (ColE8) and the focal strain (ColE2 *gfp*, or WT *gfp*). 5 μl of bacterial suspension per strain were then spotted next to each other onto a glass bottom Petri dish filled with 8 mL 0.8% w/v LB agar + PI, resulting in a distance of approximately 1.0 mm between the spot edges. After drying for ten minutes at room temperature, the plate was inverted to allow for immediate imaging of the colonies through air. Time-lapse confocal imaging of the focal strain colony edge facing the competitor was then carried out using a Zeiss EC Epiplan-Neofluar 50x air objective (NA = 0.8) on a Zeiss LSM880 confocal laser scanning unit in *Airyscan* mode, using ZEN Black software (version 14.0.18.201). Three-dimensional z-stacks at 0.36 nm intervals were then acquired every hour for eight hours. Airyscan-processed images (processing strength = 6.0) were rendered in 3D using ZEN Blue (version 2.3.69.1018) and analysed using *BiofilmQ* v0.1.4 [48]. For the data presented in Figures 4 and S2, three or four biological replicates were carried out for each strain, and for each replicate six fields of view at different distances from the colony edge were recorded at each time point. Self-lysis in each field of view was quantified by determining the volume of biomass exhibiting PI-specific fluorescence (indicating self-lysis in strain ColE2 *gfp*) relative to the total biomass. Total biomass was calculated by determining the volume of biomass exhibiting GFP-specific fluorescence (indicating either live cells or dead cells that did not self-lyse) plus biomass of those exhibiting PI-specific fluorescence.

### Statistical analysis

Statistical analysis and data visualization were performed using RStudio version 1.1.414 [49] and packages *dplyr* [50], *Rmisc* [51], *ggplot2* [52], *cowplot* [53] and *lme4* [54]. For all statistical tests, the significance level α was set to 0.01. To test whether the frequency of PI-positive cells depended on supernatant concentrations (Figures 2D and 5A), we used non-parametric Kruskal-Wallis tests. To test whether ColE2 and wildtype cells differed in their frequencies of PI-positive cells when reacting to competitor supernatant in thin colonies (Figure 3B) and three-dimensional colonies (Figure 4B and Suppl. Figure S2), we used linear models as implemented in the *lme4* package [54].

## Acknowledgements

We thank Despoina Mavridou and Diego Gonzalez for strains and plasmids, and Michael Bentley, Rolf Kümmerli, Despoina Mavridou, Jacob Palmer, Will Smith and Sofia van Moorsel for comments on the manuscript. ETG is funded by a Postdoc Mobility Fellowship from the Swiss National Science Foundation (project no. P2ZHP3_174751 and P400PB_183878). KRF is funded by European Research Council Grant 787932 and Wellcome Trust Investigator award 209397/Z/17/Z.

## Author contributions

E.G. and K.F. designed experiments; E.G. performed experiments; E.G. performed statistical analyses, E.G. and K.F. interpreted data; E.G. and K.F. wrote the manuscript.

## Supplemental Information

Supplemental information includes one table, two figures and six movies. Raw datasets underlying all figures will be deposited on the *FigShare* data repository upon acceptance.

## Supplemental Movie Legends

**Supplementary Movie S1. Cell suicide in a single *E. coli* cell shown by two markers that capture the timing of colicin production (green) and cell lysis (magenta).** More specifically, this movie shows time-lapse epifluorescence images of *E. coli* BZB1011 ColE2 pUA66-P*colE2*::*gfp* cells exposed to a 1% dilution of sterile supernatant of a strain producing colicin E8. One cell activates the ColE2 promoter (increased GPF-specific fluorescence) and subsequently undergoes self-lysis, characterized by efflux of GFP and simultaneous influx of PI-specific fluorescence. The time-lapse covers a period of 85 minutes, with 5 minutes elapsing between each frame. Scale bar, 5 µm. Selected frames from this movie are shown in Figure 2A.

**Supplementary Movie S2. Stress response (green) and cell death in *E. coli* cells unable to self-lyse.** This movie shows time-lapse epifluorescence images of *E. coli* BZB1011 WT pUA66-P*colE2*::*gfp* cells exposed to a 1% dilution of sterile supernatant of a strain producing colicin E8. Two cells activate the ColE2 promoter (increased GPF-specific fluorescence), and then fail to divide for the remainder of the observation period. No PI-specific fluorescence can be detected, indicating an intact membrane and thus no self-lysis. The time-lapse covers a period of 6 hours and 50 minutes, with 5 minutes elapsing between each frame. Scale bar, 5 µm. Selected frames from this movie are shown in Figure 2B.

**Supplementary Movie S3. Cell suicide in several *E. coli* cells shown by two markers that capture the timing of colicin production (green) and cell lysis (magenta).** This movie shows time-lapse epifluorescence images of *E. coli* BZB1011 ColE2 pUA66-P*colE2*::*gfp* cells exposed to a 1% dilution of sterile supernatant of a strain producing colicin E8. The large majority of cells activate the ColE2 promoter (increased GPF-specific fluorescence) and subsequently undergo self-lysis, characterized by efflux of GFP and simultaneous influx of PI-specific fluorescence. The time-lapse covers a period of 4 hours, with 5 minutes elapsing between each frame. Scale bar, 10 µm. A selected frame from this movie is shown in Supplementary Figure S1A.

**Supplementary Movie S4. Stress response (green) and cell death in *E. coli* cells unable to self-lyse.** This movie shows time-lapse epifluorescence images of *E. coli* BZB1011 WT pUA66-P*colE2*::*gfp* cells exposed to a 1% dilution of sterile supernatant of a strain producing colicin E8. The large majority of cells activate the ColE2 promoter (increased GPF-specific fluorescence), and then fail to divide for the remainder of the observation period. No PI-specific fluorescence can be detected, indicating an intact membrane and thus no self-lysis. The time-lapse covers a period of 4 hours, with 5 minutes elapsing between each frame. Scale bar, 10 µm. A selected frame from this movie is shown in Supplementary Figure S1B.

**Supplementary Movie S5. Cell suicide in *E. coli* cells growing in a colony, shown by two markers that capture the timing of colicin production (green) and cell lysis (magenta).** This movie shows time-lapse epifluorescence images of *E. coli* BZB1011 ColE2 pUA66-P*colE2*::*gfp* cells growing next to a competitor strain producing colicin E8, with the competitor colony located just outside the field of view to the left. The great majority of cells activate the ColE2 promoter (increased GPF-specific fluorescence) and subsequently undergo self-lysis, characterized by efflux of GFP and simultaneous influx of PI-specific fluorescence. The time-lapse covers a period of 4 hours, with 10 minutes elapsing between each frame. Scale bar, 100 µm. A selected frame from this movie is shown in Figure 3A.

**Supplementary Movie S6. Cell suicide in *E. coli* cells growing in a colony, shown by two markers that capture total biomass (green) and cell lysis (magenta).** This movie shows time-lapse three-dimensional confocal images of *E. coli* BZB1011 ColE2 *gfp* cells growing next to a competitor strain producing colicin E8, with the competitor colony located outside the field of view. The great majority of cells undergo self-lysis, characterized by efflux of GFP and simultaneous influx of PI-specific fluorescence. The time-lapse covers a period of 8 hours, with 30 minutes elapsing between each frame. A selected frame from this movie is shown in Figure 4A. See Figure 4A for scaling information.

## REFERENCES

1. Cascales, E., Buchanan, S.K., Duche, D., Kleanthous, C., Lloubes, R., Postle, K., Riley, M., Slatin, S., and Cavard, D. (2007). Colicin Biology. Microbiol. Mol. Biol. Rev. 71, 158–229.

2. Pugsley, A.P., Goldzahl, N., and Barker, R.M. (1985). Colicin E2 production and release by Escherichia coli K12 and other Enterobacteriaceae. J. Gen. Microbiol. 131, 2673–2686.

3. Pugsley, A.P., and Rosenbusch, J.P. (1981). Release of colicin E2 from Escherichia coli. J. Bacteriol. 147, 186–192.

4. Michel-Briand, Y., and Baysse, C. (2002). The pyocins of Pseudomonas aeruginosa. Biochimie 84, 499–510.

5. Šobotník, J., Bourguignon, T., Hanus, R., Demianová, Z., Pytelková, J., Mareš, M., Foltynová, P., Preisler, J., Cvačka, J., Krasulová, J., et al. (2012). Explosive backpacks in old termite workers. Science (80-.). 337, 436.

6. Laciny, A., Zettel, H., Kopchinskiy, A., Pretzer, C., Pal, A., Salim, K.A., Rahimi, M.J., Hoenigsberger, M., Lim, L., Jaitrong, W., et al. (2018). Colobopsis explodens sp. n., model species for studies on “exploding ants” (Hymenoptera, Formicidae), with biological notes and first illustrations of males of the Colobopsis cylindrica group. Zookeys 2018, 1–40.

7. Nouvian, M., Reinhard, J., and Giurfa, M. (2016). The defensive response of the honeybee Apis mellifera. J. Exp. Biol. 219, 3505–3517.

8. Shorter, J.R., and Rueppell, O. (2012). A review on self-destructive defense behaviors in social insects. Insectes Soc. 59, 1–10.

9. Mavridou, D.A.I., Gonzalez, D., Kim, W., West, S.A., and Foster, K.R. (2018). Bacteria Use Collective Behavior to Generate Diverse Combat Strategies. Curr. Biol. 28, 345–355.e4.

10. Mulec, J., Podlesek, Z., Mrak, P., Kopitar, A., Ihan, A., and Žgur-Bertok, D. (2003). A cka-gfp transcriptional fusion reveals that the Colicin K activity gene is induced in only 3 percent of the population. J. Bacteriol. 185, 654–659.

11. Kamenšek, S., Podlesek, Z., Gillor, O., and Žgur-Bertok, D. (2010). Genes regulated by the Escherichia coli SOS repressor LexA exhibit heterogenous expression. BMC Microbiol. 10, 283.

12. Bayramoglu, B., Toubiana, D., Van Vliet, S., Inglis, R.F., Shnerb, N., and Gillor, O. (2017). Bet-hedging in bacteriocin producing Escherichia coli populations: The single cell perspective. Sci. Rep. 7, 1–10.

13. Ozeki, H., Stocker, B.A.D., and De Margerie, H. (1959). Production of colicine by single bacteria. Nature 184, 337–339.

14. Ghazaryan, L., Tonoyan, L., Ashhab, A. Al, Soares, M.I.M., and Gillor, O. (2014). The role of stress in colicin regulation. Arch. Microbiol. 196, 753–764.

15. Ghazaryan, L., Soares, M.I.M., and Gillor, O. (2014). Auto-regulation of DNA degrading bacteriocins: Molecular and ecological aspects. Antonie van Leeuwenhoek, Int. J. Gen. Mol. Microbiol. 105, 823–834.

16. Gillor, O., Vriezen, J.A.C., and Riley, M.A. (2008). The role of SOS boxes in enteric bacteriocin regulation. Microbiology 154, 1783–1792.

17. Žgur-Bertok, D. (2012). Regulating colicin synthesis to cope with stress and lethality of colicin production. Biochem. Soc. Trans. 40, 1507–11.

18. Mader, A., Von Bronk, B., Ewald, B., Kesel, S., Schnetz, K., Frey, E., and Opitz, M. (2015). Amount of colicin release in escherichia coli is regulated by lysis gene expression of the colicin E2 operon. PLoS One 10, 1–17.

19. Pugsley, A.P., and Schwartz, M. (1983). A genetic approach to the study of mitomycin-induced lysis of Escherichia coli K-12 strains which produce colicin E2. MGG Mol. Gen. Genet. 190, 366–372.

20. Gonzalez, D., Sabnis, A., Foster, K.R., and Mavridou, D.A.I. (2018). Costs and benefits of provocation in bacterial warfare. Proc. Natl. Acad. Sci. U. S. A. 115, 7593–7598.

21. Cornforth, D.M., and Foster, K.R. (2013). Competition sensing: The social side of bacterial stress responses. Nat. Rev. Microbiol. 11, 285–293.

22. Kuhar, I., and Žgur-Bertok, D. (1999). Transcription regulation of the colicin K cka gene reveals induction of colicin synthesis by differential responses to environmental signals. J. Bacteriol. 181, 7373–7380.

23. Majeed, H., Gillor, O., Kerr, B., and Riley, M.A. (2011). Competitive interactions in Escherichia coli populations: The role of bacteriocins. ISME J. 5, 71–81.

24. Pugsley, A.P., and Schwartz, M. (1984). Colicin E2 release: lysis, leakage or secretion? Possible role of a phospholipase. EMBO J. 3, 2393–2397.

25. Götz, A., Lechner, M., Mader, A., Von Bronk, B., Frey, E., and Opitz, M. (2018). CsrA and its regulators control the time-point of Colicin E2 release in Escherichia coli. Sci. Rep. 8, 6537.

26. Yang, T.Y., Sung, Y.M., Lei, G.S., Romeo, T., and Chak, K.F. (2010). Posttranscriptional repression of the cel gene of the ColE7 operon by the RNA-binding protein CsrA of Escherichia coli. Nucleic Acids Res. 38, 3936–3951.

27. Chang, H.W., Yang, T.Y., Lei, G.S., and Chak, K.F. (2013). A novel endogenous induction of ColE7 expression in a csrA mutant of escherichia coli. Curr. Microbiol. 66, 392–397.

28. Lechner, M., Schwarz, M., Opitz, M., and Frey, E. (2016). Hierarchical Post-transcriptional Regulation of Colicin E2 Expression in Escherichia coli. PLoS Comput. Biol. 12, 1–20.

29. Berney, M., Hammes, F., Bosshard, F., Weilenmann, H.U., and Egli, T. (2007). Assessment and interpretation of bacterial viability by using the LIVE/DEAD BacLight kit in combination with flow cytometry. Appl. Environ. Microbiol. 73, 3283–3290.

30. Nadell, C.D., Drescher, K., and Foster, K.R. (2016). Spatial structure, cooperation and competition in biofilms. Nat. Rev. Microbiol. 14, 589–600.

31. Ghoul, M., and Mitri, S. (2016). The Ecology and Evolution of Microbial Competition. Trends Microbiol. 24, 833–845.

32. Gonzalez, D., Sabnis, A., Foster, K.R., and Mavridou, D.A.I. (2018). Costs and benefits of provocation in bacterial warfare. Proc. Natl. Acad. Sci. 115, 7593–7598.

33. Flemming, H.C., and Wuertz, S. (2019). Bacteria and archaea on Earth and their abundance in biofilms. Nat. Rev. Microbiol. 17, 247–260.

34. Mark Welch, J.L., Hasegawa, Y., McNulty, N.P., Gordon, J.I., and Borisy, G.G. (2017). Spatial organization of a model 15-member human gut microbiota established in gnotobiotic mice. Proc. Natl. Acad. Sci. 114, 201711596.

35. Tropini, C., Earle, K.A., Huang, K.C., and Sonnenburg, J.L. (2017). The Gut Microbiome: Connecting Spatial Organization to Function. Cell Host Microbe 21, 433–442.

36. Pugsley, A.P. (1983). Autoinduced synthesis of colicin E2. Mol. Gen. Genet. 190, 379–383.

37. Hamilton, W.D. (1964). The genetical evolution of social behaviour. J. Theor. Biol. 7, 1–16.

38. Granato, E.T., Foster, K.R., Meiller-Legrand, T.A., and Foster, K.R. (2019). The evolution and ecology of bacterial warfare. Curr. Biol. 29, 1–39.

39. Gardner, A., West, S.A., and Buckling, A. (2004). Bacteriocins, spite and virulence. Proc. R. Soc. B Biol. Sci. 271, 1529–1535.

40. Finnegan, J., and Sherratt, D. (1982). Plasmid ColE1 conjugal mobility: The nature of bom, a region required in cis for transfer. MGG Mol. Gen. Genet. 185, 344–351.

41. Fineran, P.C., Blower, T.R., Foulds, I.J., Humphreys, D.P., Lilley, K.S., and Salmond, G.P.C. (2009). The phage abortive infection system, ToxIN, functions as a protein-RNA toxin-antitoxin pair. Proc. Natl. Acad. Sci. U. S. A. 106, 894–899.

42. Smith, H.S., Pizer, L.I., Pylkas, L., and Lederberg, S. (1969). Abortive Infection of Shigella dysenteriae P2 by T2 Bacteriophage. J. Virol. 4, 162–168.

43. Chopin, M.C., Chopin, A., and Bidnenko, E. (2005). Phage abortive infection in lactococci: Variations on a theme. Curr. Opin. Microbiol. 8, 473–479.

44. Refardt, D., Bergmiller, T., and Kümmerli, R. (2013). Altruism can evolve when relatedness is low: Evidence from bacteria committing suicide upon phage infection. Proc. R. Soc. B Biol. Sci. 280.

45. Mitri, S., and Foster, K.R. (2013). The Genotypic View of Social Interactions in Microbial Communities. Annu. Rev. Genet. 47, 247–273.

46. Schindelin, J., Arganda-Carreras, I., Frise, E., Kaynig, V., Longair, M., Pietzsch, T., Preibisch, S., Rueden, C., Saalfeld, S., Schmid, B., et al. (2012). Fiji: An open-source platform for biological-image analysis. Nat. Methods 9, 676–682.

47. Meacock, O. (2020). FAST 0.9.1. Available at: 10.5281/zenodo.3630642.

48. Hartmann, R., Jeckel, H., Jelli, E., Singh, P.K., Vaidya, S., Bayer, M., Vidakovic, L., Díaz-Pascual, F., Fong, J.C.N., Dragoš, A., et al. (2019). BiofilmQ, a software tool for quantitative image analysis of microbial biofilm communities. bioRxiv, 735423.

49. R Development Core Team (2018). R: A language and environment for statistical computing. (Vienna, Austria: R Foundation for Statistical Computing).

50. Hadley Wickham, Romain François, L.H. and K.M. (2018). dplyr: A Grammar of Data Manipulation. R package version 0.7.6.

51. Hope, R.M. (2013). Rmisc: Ryan Miscellaneous. R package version 1.5.

52. Valero-Mora, P.M. (2010). ggplot2: Elegant Graphics for Data Analysis. J. Stat. Softw. 35, 212.

53. Wilke, C.O. (2019). cowplot: Streamlined Plot Theme and Plot Annotations for “ggplot2”. R package version 0.9.4.

54. Bates, D., Mächler, M., Bolker, B., and Walker, S. (2014). Fitting Linear Mixed-Effects Models using lme4.

55. Moffett, M. (2013). Comparative canopy biology and the structure of ecosystems. In Treetops at Risk: Challenges of Global Canopy Ecology and Conservation., M. Lowman, S. Devy, and T. Ganesh, eds. (Springer New York), pp. 13–54.

