## Supplementary Material for "The Evolution of Mass Cell Suicide in Bacterial Warfare"

*Table S1. Strains and plasmids used in this study*

| Name | Description | Source |
|---|---|---|
| BZB1011 WT (wildtype) | W3110, gyrA, $Str^R$ | [1] |
| WT *gfp* | BZB1011 Tn7::P*max*::*gfp* | [2] |
| ColE2 *gfp* | BZB1011 Tn7::P*max*::*gfp* pColE2 | [2] |
| WT pUA66-P*colE2*::*gfp* | BZB1011 pUA66-P*colE2*::*gfp*, $Kan^R$ | This study |
| ColE2 pUA66-P*colE2*::*gfp* | BZB1011 pColE2 pUA66-P*colE2*::*gfp*, $Kan^R$ | [2] |
| ColE1 | BZB1011 pColE1 | [3] |
| ColE8 | BZB1011 pColE8 | [2] |

*Table 2. Plasmids used in this study*

| Name | Description | Source |
|---|---|---|
| pColE1 (pColE1-K53) | Colicin E1 natural plasmid | [4] |
| pColE2 (pColE2-P9) | Colicin E2 natural plasmid | [4] |
| pColE8 (pColE8-J) | Colicin E8 natural plasmid | [4] |
| pUA66-P*colE2*::*gfp* | GFPmut3 transcribed from the *colE2* promoter, $Kan^R$ | [2] |

**Supplementary References**


1. Pugsley, A.P., and Schwartz, M. (1983). A genetic approach to the study of mitomycin-induced lysis of Escherichia coli K-12 strains which produce colicin E2. MGG Mol. Gen. Genet. *190*, 366–372.
2. Mavridou, D.A.I., Gonzalez, D., Kim, W., West, S.A., and Foster, K.R. (2018). Bacteria Use Collective Behavior to Generate Diverse Combat Strategies. Curr. Biol. *28*, 345-355.e4.
3. Gonzalez, D., Sabnis, A., Foster, K.R., and Mavridou, D.A.I. (2018). Costs and benefits of provocation in bacterial warfare. Proc. Natl. Acad. Sci. U. S. A. *115*, 7593–7598.
4. Ghazaryan, L., Tonoyan, L., Ashhab, A. Al, Soares, M.I.M., and Gillor, O. (2014). The role of stress in colicin regulation. Arch. Microbiol. *196*, 753–764.

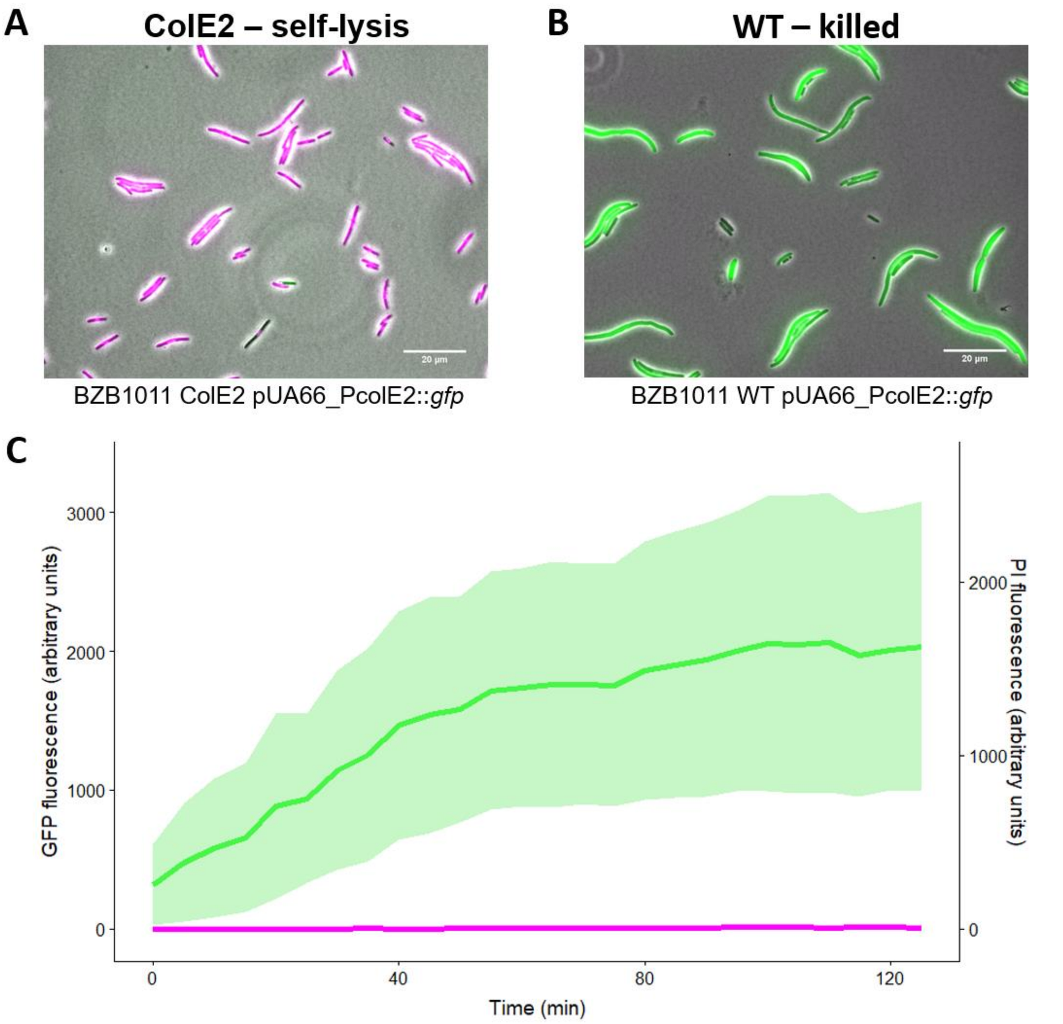


**Figure S1. *E. coli* cells unable to self-lyse do not exhibit elevated PI-fluorescence. (A and B)** Representative images of ColE2 pUA66-P*colE2*::*gfp* cells having undergone self-lysis **(A)** or BZB1011 WT pUA66-P*colE2*::*gfp* cells having been killed by a foreign DNase colicin **(B)**. Phase-contrast channel, GFP channel and propidium iodide (PI) channel are overlaid in all images. GFP signal indicates colicin promoter activation. PI signal indicates membrane permeabilization, i.e. self-lysis. Absence of PI signal in a non-dividing, dead cell is indicative of an intact membrane and hence killing by the action of the foreign DNase colicin. Image (A) represents the final timepoint of the dataset shown in Figure 2C. Image (B) represents the final timepoint of the dataset shown in (C) here. Scale bars, 20 µm. **(C)** Fluorescence signals in WT cells responding to colicin E8. WT pUA66-P*colE2*::*gfp* cells were exposed to a 1% dilution of supernatant of a colicin E8-producing strain and imaged for up to six hours. Individual cell fluorescence tracks are shown for the GFP channel (green) and PI channel (magenta). Thick lines and shaded areas indicate the mean and standard deviation across n = 31 tracked cells in the same field of view. See Movie S4.

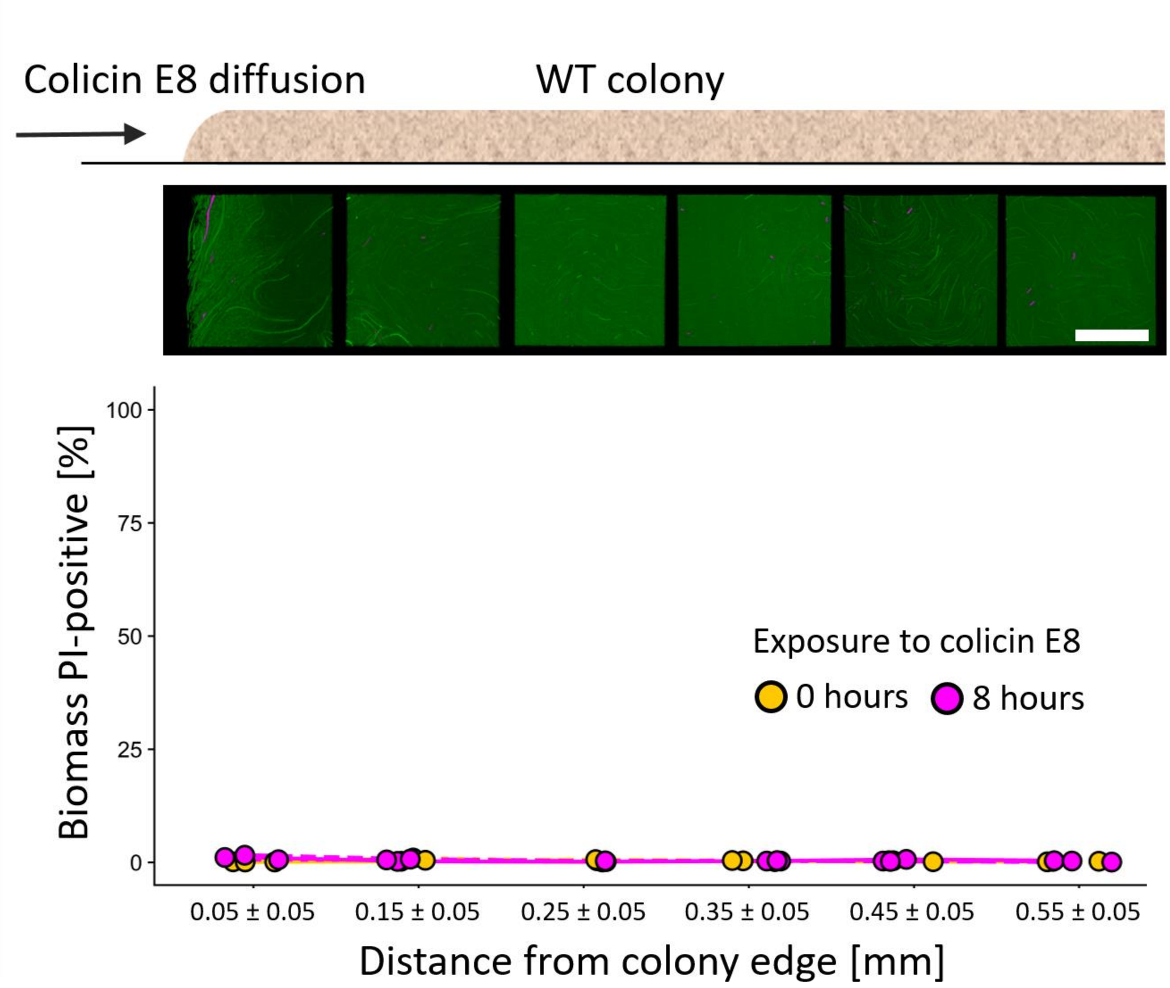


**Figure S2. Three-dimensional *E. coli* colonies unable to self-lyse do not exhibit elevated PI-fluorescence.** Quantification of PI-specific fluorescence in three-dimensional colonies exposed to colicin E8 produced by a nearby colony. Cells of a focal strain producing GFP constitutively but not capable of producing colicin E2 or self-lysing (WT *gfp*) were grown in a three-dimensional colony on nutrient medium supplement with propidium iodide (PI, 1µg/mL) next to a strain producing colicin E8. The focal colony was imaged for 8 hours using time-lapse 3D confocal microscopy at six locations situated at different distances from the colony edge facing the competitor. The proportion of PI-specific fluorescence at each location was quantified by determining the volume of biomass exhibiting PI-specific fluorescence relative to the total biomass, after zero or eight hours of exposure to colicin E8. Total biomass was calculated by determining the volume of biomass exhibiting GFP-specific fluorescence (indicating either live cells or dead cells) plus biomass of those exhibiting PI-specific fluorescence. Line-types indicate three independent biological replicates. We did not detect a statistically significant increase in PI-specific fluorescence over the observation period at any location (linear model: $F(1,4) \leq 11.89$ 212.2; $p > .01$). Images above each distance point show 3D renderings of confocal images of the same replicate colony acquired at the respective position after eight hours of exposure, viewed from above. Scale bar, 50 µm.
